# A data-driven interactome of synergistic genes improves network based cancer outcome prediction

**DOI:** 10.1101/349688

**Authors:** Amin Allahyar, Joske Ubels, Jeroen de Ridder

## Abstract

Robustly predicting outcome for cancer patients from gene expression is an important challenge on the road to better personalized treatment. Network-based outcome predictors (NOPs), which considers the cellular wiring diagram in the classification, hold much promise to improve performance, stability and interpretability of identified marker genes. Problematically, reports on the efficacy of NOPs are conflicting and for instance suggest that utilizing random networks performs on par to networks that describe biologically relevant interactions. In this paper we turn the prediction problem around: instead of using a given biological network in the NOP, we aim to identify the network of genes that truly improves outcome prediction. To this end, we propose SyNet, a gene network constructed ab initio from synergistic gene pairs derived from survival-labelled gene expression data. To obtain SyNet, we evaluate synergy for all 69 million pairwise combinations of genes resulting in a network that is specific to the dataset and phenotype under study and can be used to in a NOP model. We evaluated SyNet and 11 other networks on a compendium dataset of >4000 survival-labelled breast cancer samples. For this purpose, we used cross-study validation which more closely emulates real world application of these outcome predictors. We find that SyNet is the only network that truly improves performance, stability and interpretability in several existing NOPs. We show that SyNet overlaps significantly with existing gene networks, and can be confidently predicted (~85% AUC) from graph-topological descriptions of these networks, in particular the breast tissue-specific network. Due to its data-driven nature, SyNet is not biased to well-studied genes and thus facilitates post-hoc interpretation. We find that SyNet is highly enriched for known breast cancer genes and genes related to e.g. histological grade and tamoxifen resistance, suggestive of a role in determining breast cancer outcome.

**Author Summary:** Cancer is caused by disrupted activity of several pathways. Therefore, outcome predictors analyze patient’s expression profiles from perspective of gene groups collected from interactomes (e.g. protein interaction networks). These Network based Outcome Predictors (NOPs) hold potential to facilitate identification of dysregulated pathways and delivering improved prognosis. Nonetheless, recent studies revealed that compared to classical models, neither performance nor consistency can be improved using NOPs.

We argue that NOPs can only perform well under guidance of suitable networks. The commonly used networks may miss associations specially for under-studied genes. Additionally, these networks are often generic with low resemblance to perturbations that arise in cancer.

To address this issue, we exploit ~4100 samples and infer a disease specific network called SyNet linking synergistic gene pairs that collectively show predictivity beyond individual performance of genes.

Using identical datasets, we show that a NOP yields superior performance merely by considering groups of genes in SyNet. Further, NOP performance severely reduces if SyNet nodes are shuffled, confirming relevance of SyNet links.

Due to simplicity of our approach, this framework can be used for any phenotype of interest. Our findings represent the value of network-based models and crucial role of interactome in their performance.

## 1 Introduction

Metastases at distant sites (e.g. in bone, lung, liver and brain) is the major cause of death in breast cancer patients [1]. However, it is currently difficult to assess tumor progression in these patients using common clinical variables (e.g. tumor size, lymph-node status, etc.) [2]. Therefore, for 80% of these patients chemotherapy is prescribed [3]. Meanwhile, randomized clinical trials showed that at least 40% of these patients survive without chemotherapy and thus unnecessarily have to suffer from the toxic side effect of this treatment [3,4]. For this reason, substantial efforts have been made to derive molecular classifiers that can predict clinical outcome based on gene expression profiles obtained from the primary tumor at the time of diagnosis [5,6].

An important shortcoming in molecular classification is that ‘cross-study’ generalization is often poor [7]. This means that prediction performance decreases dramatically when a classifier trained on one patient cohort is applied to another one [8]. Moreover, the gene signatures found by these classifiers vary greatly, often sharing only few or no genes at all [9–11]. This lack of consistency casts doubt on whether the signatures capture true ‘driver’ mechanisms of the disease or rather subsidiary ‘passenger’ effects [12].

Several reasons for this lack of consistency have been proposed, including small sample size [11,13,14], inherent measurement noise [15] and batch effects [16,17]. Apart from these technical explanations, it is recognized that traditional models ignore the fact that genes are organized in pathways [18]. One important cancer hallmark is that perturbation of these pathways may be caused by deregulation of disparate sets of genes which in turn complicates marker gene discovery [19,20].

To alleviate these limitations, the classical models are superseded by Network-based Outcome Prediction (NOP) methods which incorporate gene interactions in the prediction model [21]. NOPs have two fundamental components: aggregation and prediction. In the aggregation step, genes that interact, belong to the same pathway or otherwise share functional relation are aggregated (typically by averaging expressions) into so called “meta-genes” [22]. This step is guided by a secondary data source describing gene-gene interactions such as cellular pathway maps or protein-protein interaction networks. In the consequent prediction step, meta-genes are selected and combined into a trained classifier, similar to a traditional classification approach. Several NOPs have been reported to exhibit improved discriminative power, enhanced stability of the classification performance and signature and better representation of underlying driving mechanisms of the disease [18,23–25].

In recent years, a range of improvements to the original NOP formulation has been proposed. In the prediction step, various linear and nonlinear classifiers have been evaluated [26,27]. Problematically, the reported accuracies are often an overestimation as many studies neglected to use cross-study evaluation scheme which more closely resembles the real world application of these models [7]. Also for the aggregation step which is responsible for forming meta-genes from gene sets several distinct approaches are proposed, such as clustering [23] and greedy expansion of seed genes into subnetworks [18]. Moreover, in addition to simple averaging, alternative means by which genes can be aggregated, such as linear or nonlinear embeddings, have been proposed [17,28]. Most recent work combines these steps into a unified model [8,29].

Despite these efforts and initial positive findings, there is still much debate over the utility of NOPs compared to classical methods, with several studies showing no performance improvement [21,30,31]. Perhaps even more striking is the finding that utilizing a permuted network [31] or aggregating random genes [10] performs on par with networks describing true biological relationships. Several meta-analyses attempting to establish the utility of NOPs have appeared with contradicting conclusions. Notably, Staiger et al. compared performance of nearest mean classifier [32] in this setting and concluded that network derived meta-genes are not more predictive than individual genes [21,31]. This is in contradiction to Roy et al. who achieved improvements in outcome prediction when genes were ranked according to their t-test statistics compared to their pagerank property [33] in PPI network [28,34]. It is thus still an open question whether NOPs truly improve outcome prediction in terms of predictive performance, cross-study robustness or interpretability of the gene signatures.

A critical - yet often neglected - aspect in the successful application of NOPs is the contribution of the biological network. In this regard, it should be recognized that many network links are unreliable [35,36], missing [37] or redundant [38] and considerable efforts are being made to refine these networks [37,39–41]. In addition, many links in these networks are experimentally obtained from model organisms and therefore may not be functional in human cells [42–44]. Finally, most biological networks capture only a part of a cell’s multifaceted system [45]. This incomplete perspective may not be sufficient to link the wide range of aberrations that may occur in a complex and heterogeneous disease such as breast cancer [46,47]. Taken together, these issues raise concerns regarding the extent to which the outcome predictors may benefit from inclusion of common biological networks in their models.

In this work, we propose to construct a network *ab initio* that is specifically designed to improve outcome prediction in terms of cross-study generalization and performance stability. To achieve this, we will effectively turn the problem around: instead of using a given biological network, we aim to use the labelled gene expression datasets to identify the network of genes that truly improves outcome prediction (see **Figure 1** for a schematic overview).

**Figure 1.**
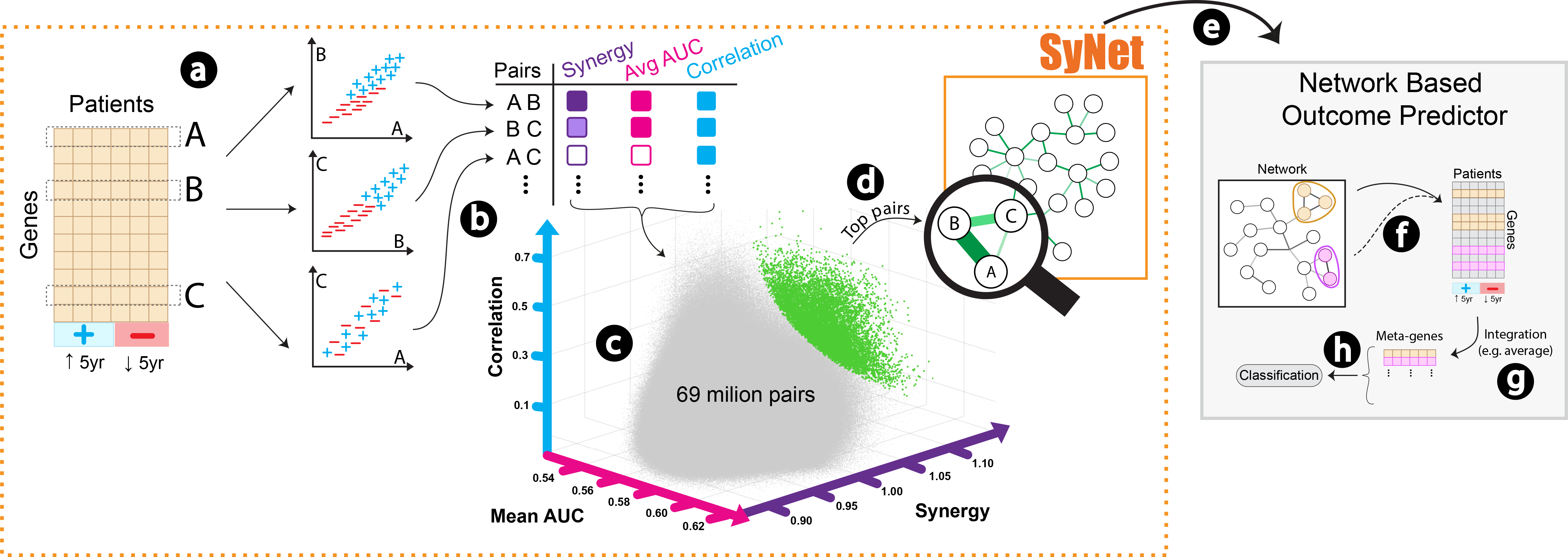
Schematic overview of SyNet inference and NOP training. For every 69 million combinations of gene pairs **(a)** we compute three criteria including synergy (*S*_*ij*_, purple), average AUC (*M*_*ij*_, pink), and correlation (*C*_*ij*_, blue) **(b)**. These three criteria form a threedimensional space **(c)** from which Fitness (*F*_*ij*_) can be calculated for each pair. Top pairs (green dots) in this space collectively form SyNet **(d)**. SyNet is subsequently used in a NOP **(e)**, in which the links in SyNet guide the construction of “meta-genes”. Within a NOP groups of genes are formed **(f)** and then integrated into meta-genes (typically using averaging) **(g)**. The constructed meta-genes are then used as regular features to train standard classifiers **(h)**. The phenotype of interest is patient outcome (i.e. 5-year survival).

Our approach relies on the identification of *synergistic gene pairs*, i.e. genes whose joint prediction power is beyond what is attainable by both genes individually [48]. To identify these pairs, we employed grid computing to evaluate all 69 million pairwise combinations of genes. The resulting network, called SyNet, is specific to the dataset and phenotype under study and can be used to successfully infer a NOP model.

To obtain SyNet, and allow for rigorous cross-study validation, a dataset of substantial size is required. For this reason, we combined 14 publicly available datasets to form a compendium encompassing 4129 samples. To the best of our knowledge, the data combined in this study represents the largest breast cancer gene expression compendium to date. Further, to ensure unbiased evaluation, sample assignments in the inner as well as the outer cross-validations folds are kept equal across all assessments throughout the paper.

In the remainder of this paper, we will demonstrate that integrating genes based on SyNet provides superior performance and stability of predictions when these models are tested on independent cohorts. In contrast to previous reports, where shuffled versions of networks also performed well, we show that the performance drops substantially when SyNet links are shuffled, suggesting its connections are truly informative. We further evaluate the content and structure of SyNet by overlaying it with known genesets and existing networks, revealing marked enrichment for known breast cancer prognostic markers. While overlap with existing networks is highly significant, the majority of direct links in SyNet is absent from these networks. Interestingly, they can be reliably predicted from existing networks when more complex topological descriptions are taken into account. Taken together, our findings suggest that compared to generic gene networks, phenotype specific networks, which are derived directly from labeled data, can provide superior performance while at the same time revealing valuable insight into etiology of breast cancer.

## 2 Results

### 2.1 SyNet improves NOP performance

We first evaluated NOP performance for three existing methods (Park, Chuang and Taylor) and the GL when supplied with a range of networks, including generic networks, tissue specific networks and SyNet. The AUC of the four NOPs, presented in **Figure 2**, clearly demonstrates that SyNet improves the performance of all NOPs, except for the Park method in which performs on par using the Corr network. None of the existing NOPs truly outperform the classifier Lasso (i.e. baseline; rightmost bar in **Figure 2**) which is the top performing regular classifier in our evaluations; see **S3** for details).

**Figure 2.**
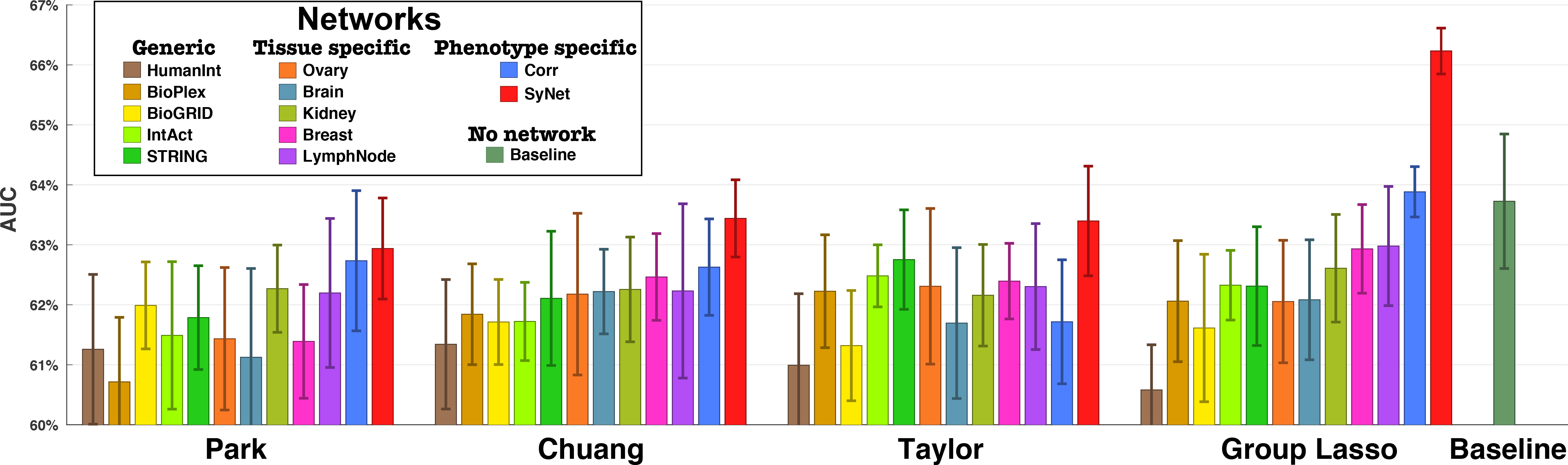
Performance comparison of NOPs for 4 methods and 12 networks including SyNet. Each bar represents average performance in terms of the AUC. The rightmost bar represents the performance of standard Lasso which considers all individual genes as features (i.e. no network). Error bars represent the standard deviation across 10 repeats.

The GL clearly outperforms all other methods, in particular when it exploits the information contained in SyNet. This corroborates our previous finding [8] that existing methods which construct meta-genes by averaging are suboptimal (see **S1** for a more extensive analysis). The GL using the Corr network also outperforms the baseline model, albeit non-significantly (p~0.6), which is in line with previous reports [23]. It should be noted that across all these experiments an identical set of samples is used to train the models so that any performance deviation must be due to differences in (i) the set of utilized genes or (ii) the integration of the genes into metagenes. In the next two sections, we will investigate these factors in more details.

### 2.2 SyNet provides feature selection capabilities

Networks only include genes that are linked to at least one other gene. As a result, networks can provide a way of ranking genes based on the number and weight of their connections. One explanation for why NOPs can outperform regular classifiers is that networks provide an a priori gene (feature) selection [31]. To test this hypothesis and determine the feature selection capabilities of SyNet, we compare classification performances obtained using the baseline classifier (i.e. Lasso) that is trained using enclosed genes in each network. While this classifier performs well (see **S3** for details), it cannot exploit information contained in the links of given network. So any performance difference must be due to the genes in the network. The number of genes in the network was varied by thresholding the weighted edges in the network and removing unconnected genes. The edge weight threshold and the Lasso regularization parameter were determined using a cross-validated grid search (see **S5** for details). **Figure 3** provides the optimal performances for 12 distinct networks along with number of genes used in the final model (i.e. genes with non-zero Lasso coefficients). We also included the baseline model where all genes are utilized to train Lasso classifier (rightmost bar).

**Figure 3.**
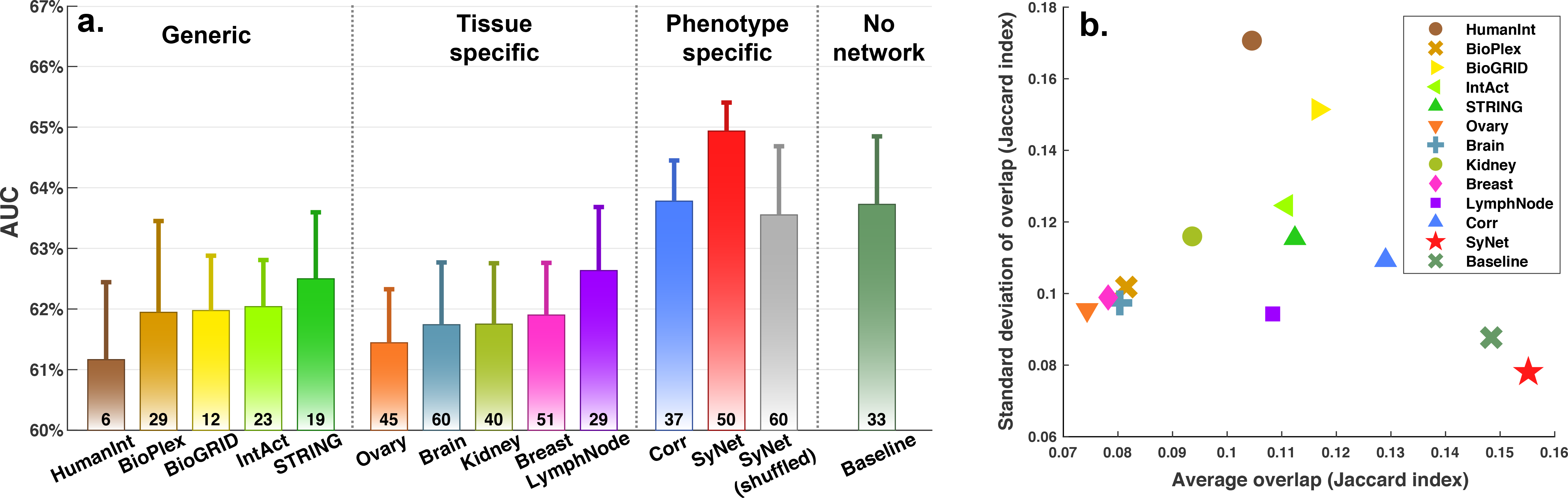
Performance comparison between networks when interconnections are ignored and genes contained in each network are utilized to train Lasso. **a.** Performance (AUC) of Lasso classification using individual genes in 12 networks. Numbers below each bar represent the median number of non-zero coefficients after training Lasso across 10 repeats and 14 folds. **b.** Stability of identified signatures measured by overlap between identified gene sets using Jaccard index. X and y-axis represent average and standard deviation of the Jaccard index measured across 10 repeats and 14 folds.

The results presented in **Figure 3.a** demonstrate that SyNet is the only network that performs significantly better than the baseline model which is trained on all genes. Interestingly, we observe that SyNet is the top performing network while utilizing a comparable number of genes to other networks. The second-best network is the Corr network. Compared to the Corr network, SyNet additionally had access to label information, making it also specific to the class label under study. It should be noted that the data on which SyNet and the Corr networks are constructed are completely independent from the validation data on which the performance is based due to our multi-way cross-validation scheme (see Methods and **S5**). We conclude that dataset specific networks, in particular SyNet which also exploits label information, provides a meaningful feature selection that is beneficial for classification performance.

Our result show that none of the tissue specific networks outperform the baseline. Despite the modest performance, it is interesting to observe that performance for these networks increases as more relevant tissues (e.g. breast and lymph node networks) are utilized in the classification. Additionally, we observe that tissue specific networks do not outperform the generic networks. This may be the result of the fact that generic networks predominantly contain broadly expressed genes with fundamental roles in cell function which may still be relevant to survival prediction. A similar observation was made for GWAS where SNPs in these widely-expressed genes can explain a substantial amount of missed heritability [59].

In addition to classifier performance, an important goal of employing NOPs is to identify stable gene signatures, that is, the same genes are selected irrespective of the study used to train the models. Gene signature stability is necessary to confirm that the identified genes are independent of dataset specific variations and are true biological drivers of the disease. To measure the signature consistency, we assessed the overlap of selected genes across all repeats and folds using the Jaccard Index. **Figure 3.b** shows that, using genes included in SyNet, Lasso identifies more similar genes across folds and studies compared to other networks. Surprisingly, despite the fact that the expression data from which SyNet is inferred changes in each classification fold, the signature stability for SyNet is markedly better than for generic or tissue specific networks, in which the genes are fixed. Therefore, our results demonstrate that synergistic genes in SyNet truly aid the classifier to robustly select signatures across independent studies.

### 2.3 SyNet connections are beneficial for NOP

The ultimate goal of employing NOPs compared to classical models that do not use network information is to improve prognosis prediction by harnessing the information contained in the links of the given network. Therefore, we next aimed to assess to what extend also connections between the genes, as captured in SyNet and other networks, can help NOPs to improve their performance beyond what is achievable using individual genes. As before, we utilized identical datasets (in terms of genes and samples) in inner and outer cross-validation loops to train all four NOPs as well as the baseline model which uses Lasso. As a result, in our comparison, the only difference between each NOP and the regular Lasso is the grouping of genes based on their connectivity in the networks.

Our results, presented in **Figure 4.a**, clearly demonstrate that SyNet is the only network to achieve superior prognostic prediction for unseen patients from an independent cohort. It outperforms all other tested networks, including relevant tissue specific networks. Moreover, we find that using SyNet also provides one of the largest gains in performance over its Lasso counterpart (**Figure 4.b**, x-axis), indicating that the links between genes in SyNet truly aid classification performance beyond what is obtained as a result of the feature selection capabilities.

**Figure 4.**
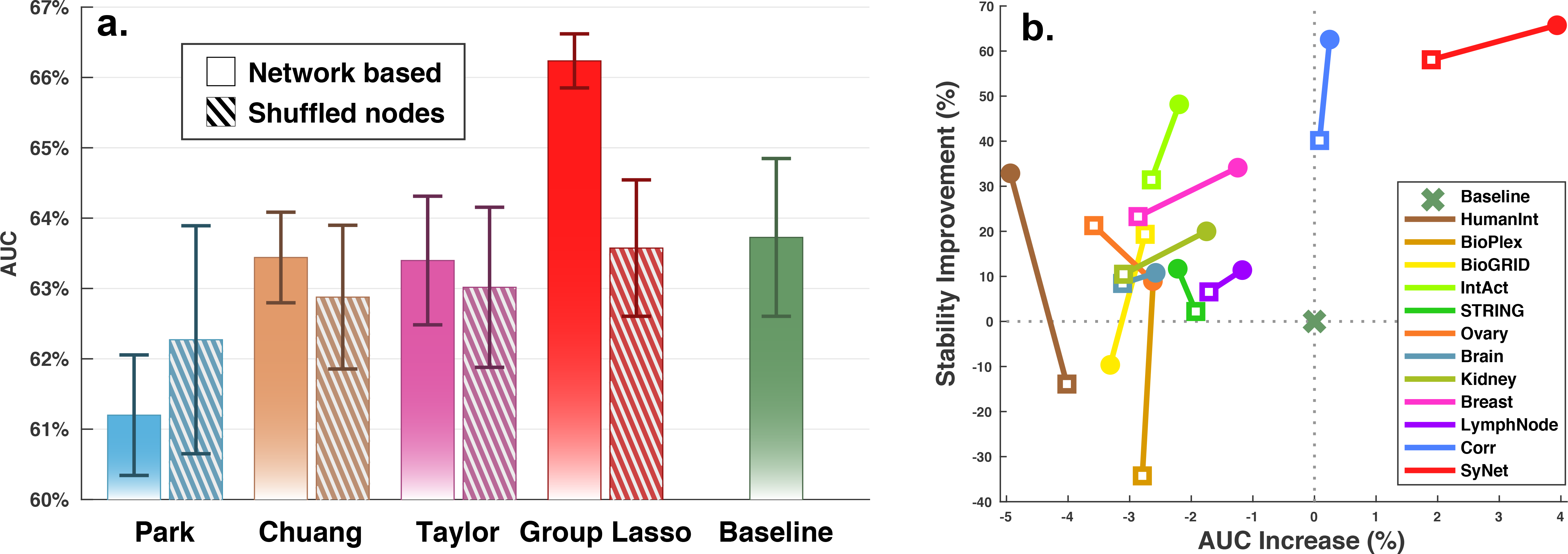
Performance of NOP models trained using SyNet compared to a shuffled version of this network. **a.** Bars indicate average performance of models across repeats and error bars denote the corresponding standard deviation. Solid bars represent average performance of models trained using SyNet. Dashed bars denote performance of the same model using randomized SyNet, i.e. the same genes are present but randomly connected. **b.** Improvement of performance (x-axis) and stability (in terms of the standard deviation of the AUC; y-axis) compared to the baseline model. Square and circle markers represent performance obtained using genes (i.e. Lasso) and the network (i.e. GL), respectively.

To confirm that NOP performance using SyNet is the result of the network structure, we also applied the GL to a shuffled version of SyNet (**Figure 4.a**). We observe a substantial deterioration of the AUC supporting the conclusion that not only the genes, but also links contained in SyNet are relevant to achieve good prediction. Moreover, this observation rules out that the GL by itself is able to provide enhanced performance compared to standard Lasso. Result of similar assessment for Corr network is given in (Figure **S12**).

Another important property of an outcome predictor is to yield comparable performance irrespective of dataset used for training the model (i.e. performance stability). This is a highly desirable quality, as concerns have been raised regarding the highly variable performances of breast-cancer classifiers applied to different cohorts [7,60]. **Figure 4.b** clearly demonstrates that a NOP model guided by SyNet not only provide superior performance, it also offers improved stability of the classification performance.

We further compared performances when each network contains equal number of links and observe a similar trend in performance (see **S7** for details). We also considered the more complex Sparse Group Lasso (SGL), which offers an additional level of regularization. No substantial difference between GL and SGL performance was found (see **S8** for details). Likewise, we did not observe substantial performance differences when the number of genes, group size and regularization parameters were simultaneously optimized in a grid search (see **S9** for details). Together, these findings can be considered as the first unbiased evidence of true classification performance improvement in terms of average AUC and classification stability by a NOP.

### 2.4 Gene enrichment analysis for SyNet

Many curated biological networks suffer from bias since genes with well-known roles are the subject of more experiments and thus get more extensively and accurately annotated [61]. Post-hoc interpretation of the features used by NOPs, often by means of an enrichment analysis, will therefore be affected by the same bias. SyNet does not suffer from such bias, as its inference is purely data driven. Moreover, since SyNet is built based on gene pairs that contribute to the prediction of clinical outcome, we expect that the genes included in the network not only relate to breast cancer; they should play a role in determining how aggressively the tumor behaves, how advanced the disease is or how well it responds to treatment.

To investigated the relevance of genes contained in SyNet in the development of breast cancer and, more importantly, clinical outcome, we ranked all pairs according to their median Fitness (*F*_*ij*_) across 14 studies and selected the top 300 genes (encompassing 3544 links). This cutoff was frequently chosen by the Lasso as the optimal number of genes in SyNet (see section 3.1). **Figure 5** depicts a visualization of this network revealing three main subnetworks and a few isolated gene pairs. We performed functional enrichment of all genes using Ingenuity Pathway Analysis (IPA) [62], and functional enrichment of the subcomponents of the three large subnetworks using Gene Set Enrichment Analysis (GSEA) [63].

**Figure 5.**
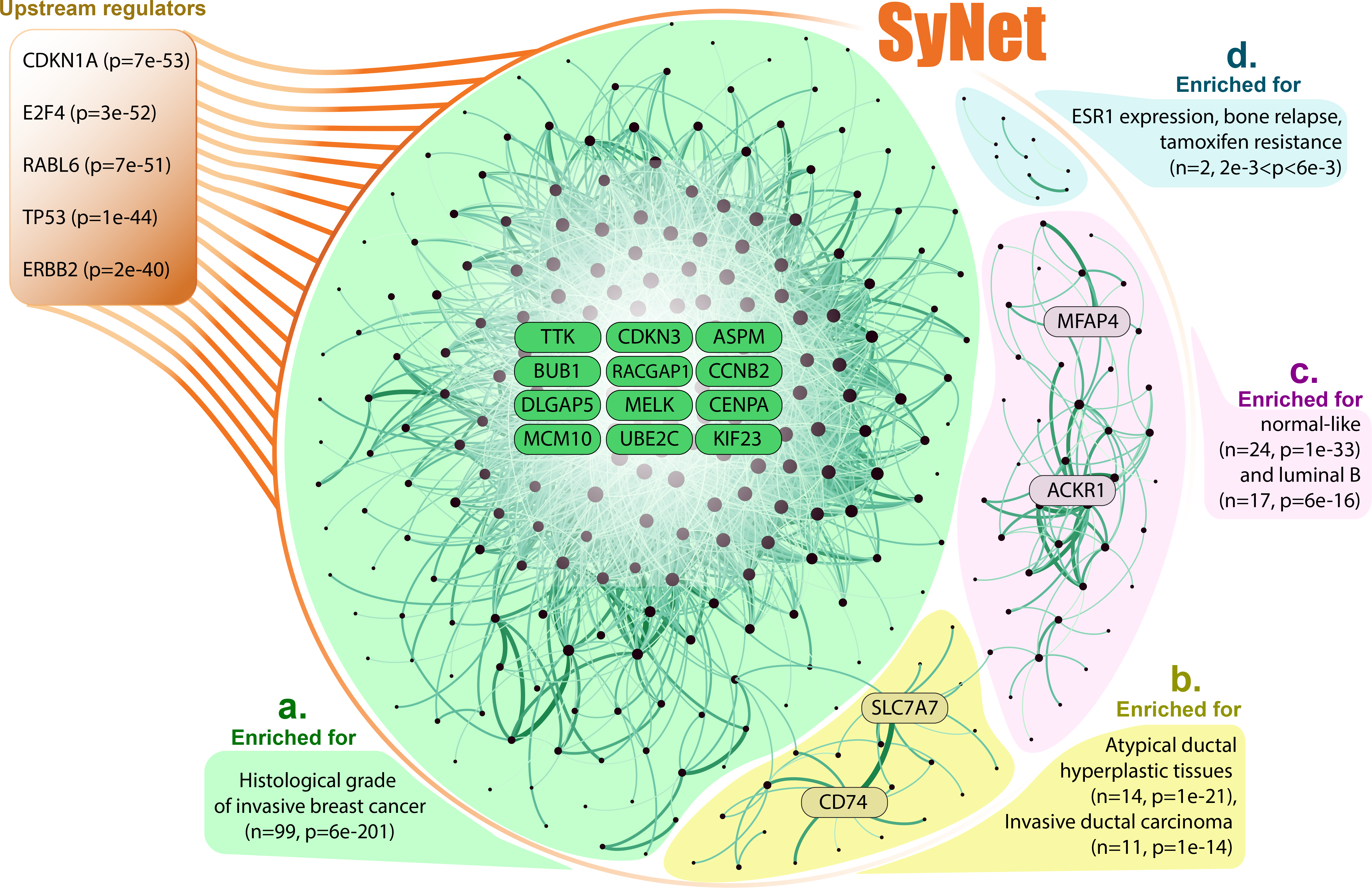
Visualization of SyNet. SyNet consists of three main subnetworks (**a**, **b** and **c**) and five separated gene pairs (**d**). Node size represents degree of node and link thickness indicates fitness of the corresponding pair. **a.** The largest subnetwork encompassing 223 genes is enriched for histologic grade of invasive breast cancer tumors. **b.** The second subnetwork is directly connected to the first cluster and contains risk factors for developing breast cancer. **c**. The third cluster is enriched for genes upregulated in normal-like subtype of breast cancer. **d.** Out of five pairs, only TFF3 and TFF1 pair is enriched for genes up-regulated in early primary breast tumors.

IPA reveals that out of 300 genes in SyNet, 287 genes have a known relation to cancer (2e-06<p<1e-34) of which 222 are related to reproductive system disease (2e-06<p<1e-34). Furthermore, the top five upstream regulators of genes in SyNet (orange box, **Figure 5**) are CDKN1A, E2F4, RABL6, TP53 and ERBB2, all of which are well known players in the development of breast cancer [64–68]. The mean degree of the 300 genes in the network build by SyNet is 24, but there are 12 genes which have a degree of 100 or above: ASPM [69], BUB1 [70], CCNB2 [71], CDKN3 [72], CENPA [73], DLGAP5 [74], KIF23 [75], MCM10 [76], MELK [77], RACGAP1 [78], TTK [79] and UBE2C [80]. All these genes play a vital role in progression through the cell cycle and mitosis, by ensuring proper DNA replication, correct formation of the mitotic spindle and proper attachment to the centromere.

In addition to a clear involvement of genes linked to breast cancer generically, we also find clear indications that the genes in SyNet are relevant to clinical outcome and prognosis of the disease. For instance, the most highly enriched cluster (green cluster **Figure 5**) is found to be associated to histological grade of the tumor. The histological grade, which is based on the morphological characteristics of the tumor, has been shown to be informative for the clinical behavior of the tumor and is one of the best-established prognostic markers [81]. Notably, the blue cluster is enriched for genes involved in tamoxifen resistance, one of the important treatments of ER-positive breast cancer.

Two other sub-clusters (yellow and purple in **Figure 5**), contain genes from distinctly different biological processes than the main cluster. In these clusters we also observe clear hub genes: SLC7A7 and CD74 in the yellow and ACKR1 and MFAP4 in the purple cluster. ACKR1 is a chemokine receptor involved in the regulation of the bio-availability of chemokine levels and MFAP4 is involved in regulating cell-cell adhesion. The recruitment of cells, as regulated by chemokines, and reducing cell-cell adhesion both play an important role in the process of metastasis. CD74 has also been linked to metastasis in triple negative breast cancer [82]. Metastasis, and not the primary tumor, is the main cause of death in breast cancer [3].

IPA highly significantly identifies the SyNet genes as upstream regulators of canonical pathways implicated in breast cancer (**Figure 5**), such as Cell Cycle Control of Chromosomal Replication (8e-18), Mitotic Roles of Polo-Like Kinase (4e-15), Role of CHK Proteins in Cell Cycle Checkpoint Control (6e-12), Estrogen-mediated S-phase Entry (2e-11), and Cell Cycle: G2/M DNA Damage Checkpoint Regulation (5e-10). Although all cancer cells deregulate cell cycle control, the degree of dysregulation may contribute to a more aggressive phenotype. For instance, it is recognized that the downregulation of certain checkpoint regulators are related to a worse prognosis in breast cancer [83,84].

In summary, SyNet predominantly appears to contain genes relevant to two main processes in the progression of breast cancer: increased cell proliferation and the process of metastasis. Although many genes have not previously been specifically linked to breast cancer prognosis, their role in regulating different stages of replication and mitosis points to a genuine biological role in the progression and prognosis of breast cancer.

### 2.5 Similarity of SyNet to existing biological networks

We next sought to investigate the similarity between SyNet and existing biological networks that directly or indirectly capture biological interactions. To enable a comparison with networks of different sizes, we compare the observed overlap (both in terms of genes as well as links) to the distribution of expected overlap obtained by shuffling each network 1000 times. Overlap is determined for varying network sizes by thresholding the link weights such that a certain percentage of genes or links remains. Results are reported in terms of a z-score in **Figure 6**.

**Figure 6.**
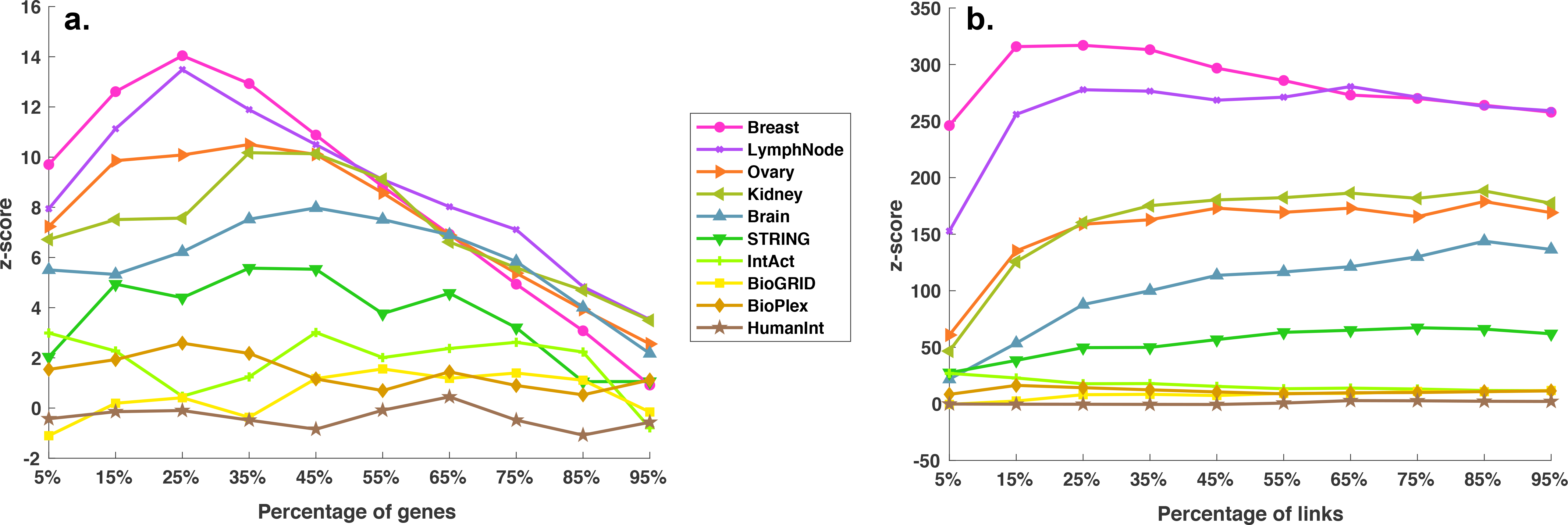
Similarity of existing biological networks to SyNet in terms of genes (**a.**) and links (**b.**). The x-axis represents the percentage of top gene/links used, the y-axis the z-score of observed vs. expected number of gene/links. The z-score is calculated by relating the observed number of SyNet gene/links that are present in existing biological networks to the expected distribution. To calculate the expected distribution, genes in biological networks are shuffled.

**Figure 6.a** shows that for the majority of networks a significantly higher than expected number of SyNet genes is contained in the top of each network. The overlap is especially pronounced for the tissue specific networks, in particular the Breast and Lymph node specific network, supporting our observation that SyNet contains links that are relevant for breast cancer. The enrichment becomes even more significant when considering the overlap between the links (**Figure 6.b**). In this respect, SyNet also is clearly most similar to the Breast and Lymph node specific networks. We confirmed that these enrichments are not only driven by the correlation component of SyNet by repeating this analysis with the correlation component removed (see **S10**). It should moreover be noted that, although a highly significant overlap is observed, the vast majority of SyNet genes and links are not present in the existing networks, explaining the improved performance obtained with NOPs using SyNet. Specifically, out of the 300 genes in SyNet, only 142 are contained within the top 25% of genes (n=1005) in the Breast specific network, and 151 in the top 25% of genes (n=1290) in the Lymph node specific network. Similarly, out of the 3544 links in SyNet, only 1182 are contained within the top 25% of links (n=12500) in the Breast specific network, and 617 in the top 25% (n=12500) of the Lymph node specific network (see **S11** for details).

### 2.6 Higher order structural similarity of SyNet and existing biological networks

In addition to direct overlap, we also aimed to investigate if genes directly connected in SyNet may be indirectly connected in existing networks. To assess this for each pair of genes in SyNet, we computed several topological measures characterizing their (indirect) connection in the biological networks. We included degree (**Figure 7.a**), shortest path (**Figure 7.b**) and jaccard (**Figure 7.c**) (see supplementary text for details). To produce an edge measure from degree and pagerank (which are node based), we computed the average degree and pagerank of genes in a pair respectively. Furthermore, we produced an expected distribution for each pair by computing the same topological measures for one of the genes and another randomly selected gene. The results from this analysis supports our previous observation that the information contained in the links of SyNet is substantially - yet partially - overlapping with the information in the existing networks. Notably, the similarity increases for networks of increased relevance to the tissue in which the gene expression data is measured (i.e. breast tissue).

**Figure 7.**
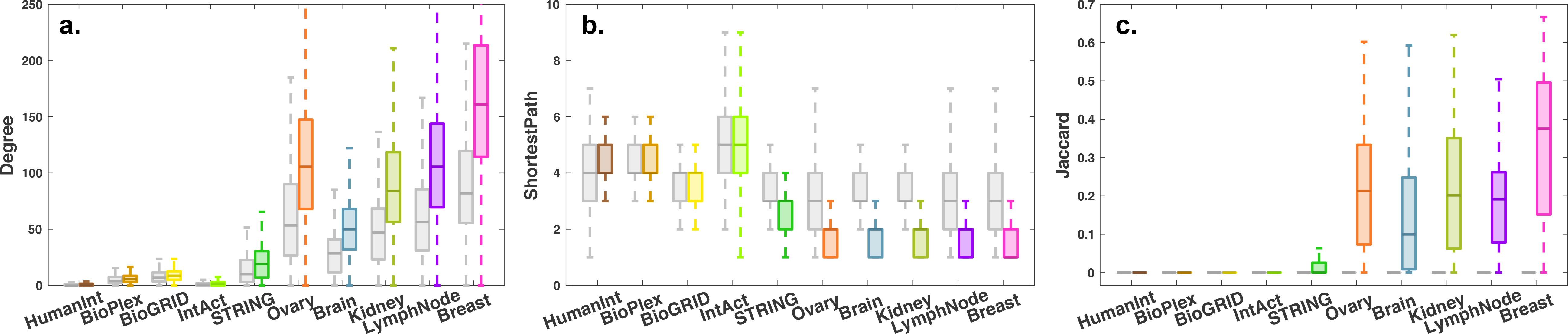
Comparison of three topological measures calculated over biological networks. Each color represents a network. Gray boxes represent the same topological measures calculated on the shuffled network.

### 2.7 Predicting SyNet links from biological networks

Encouraged by the overlap with existing biological networks, we next asked whether links in SyNet can be predicted from the complete collection of existing networks. To this end we characterized each gene-pair by a set of 12 graph-topological measures that describe local and global network structure around each gene-pair. In addition to the degree, shortest path and Jaccard, we included several additional graph-topological measures including direct link, page rank (with four betas), closeness centrality, clustering coefficient and eigenvector centrality (see supplementary text for details). While converting node-based measures to edge based measures, in addition to using the average, we also used the difference between the score for each gene in the pair, similar to our previous work [85]. We applied these measures to all 10 networks in our collection yielding a total of 210 features. The gene-pairs are labeled according to their presence or absence in SyNet. Inspection of this dataset using the t-SNE [57] reveals that the links in SyNet occupy a distinct part of the 2D embedding obtained (**Figure 8.a**).

**Figure 8.**
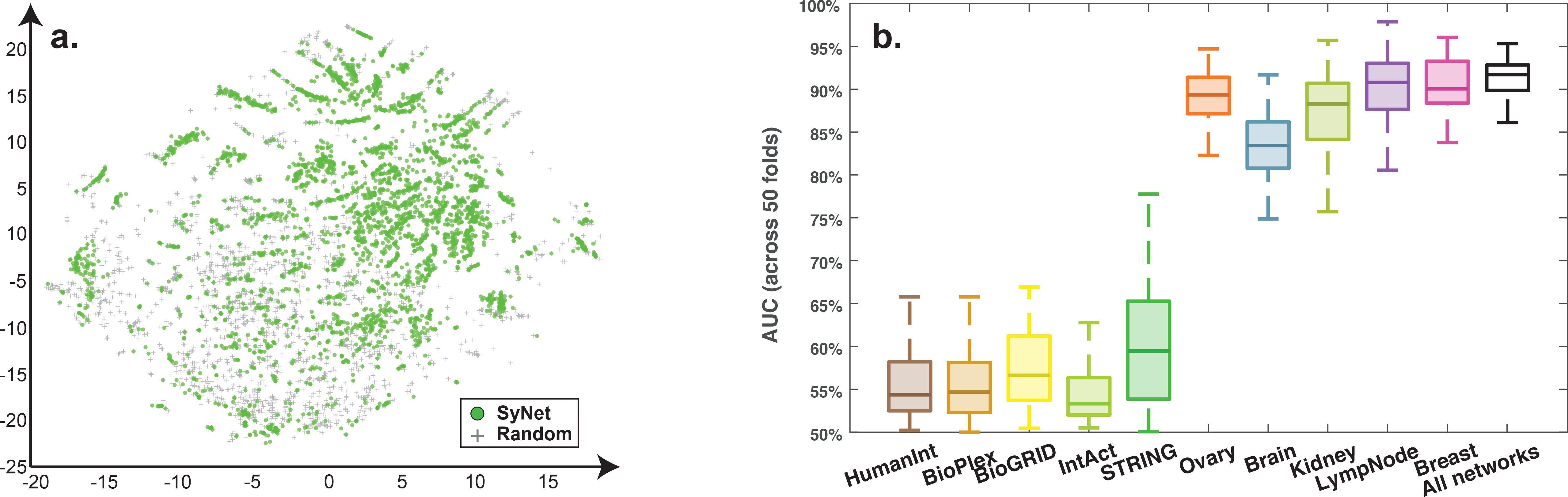
Characterizing SyNet links by a range of graph topological measures. **a.** t-SNE (unsupervised) visualization of the combined 180 topological measures. Each dot represents one gene pair. Green dots indicate SyNet links while gray markers represent an equal number of random pairs. **c.** Performance of Lasso model trained over all topological measure for different networks and all networks combined (rightmost bar).

We trained a Lasso and assessed classification performance in a 50-fold cross validation scheme where in each fold 1/50 of pairs in SyNet is kept hidden and the rest of pairs is utilized to train the classifier. To avoid information leakage in this assessment, we removed gene pairs from the training set in case one of the genes is present in the test set. Based on this analysis we find that a simple linear classifier can reach ~85% accuracy in predicting the synergistic gene relationships from SyNet (**Figure 8.b**, rightmost bar). The contribution from generic networks is markedly smaller than for the tissue specific networks. In particular the networks relevant to breast cancer are highly informative, to the extent that combining multiple networks no longer improves prediction performance. Further investigation of feature importance revealed that the page rank topological measure was commonly used as a predictive marker. Apparently, while direct overlap between SyNet and existing networks is modest, the topology of the breast specific networks is highly informative for the links contained in SyNet. This corroborates findings from Winter et al. in which the page rank topological measure was proposed to identify relevant genes in outcome prediction [33,34,86].

## 3 Discussion and Future work

Although the principle of using existing knowledge of the cellular wiring diagram to improve performance, robustness and interpretability of gene expression classifiers appears attractive, contrasting reports on the efficacy of such approach have appeared in literature [21,28,31,34]. Consensus in this field has particularly been frustrated by an evaluation of a limited set of sub-optimal classifiers [21,23,28,34], small sample size [18,24,26], or the use of standard K-fold crossvalidation instead of cross-study evaluation schemes [24,26]. For this reason, it remained unclear if network-based classification, and in particular network-based outcome prediction, is beneficial. Here, we present a rigorously cross-validated procedure to train and validate Group Lasso-based NOPs using a variety of networks, in particular also tissue specific networks, which have not been evaluated in the context of NOPs before.

Based on our analyses we conclude that none of the existing networks achieve improved performance compared to using properly regularized classifiers trained on all genes. In this work we therefore present a novel gene network, called SyNet, which is computationally derived directly from the labeled dataset. The links in SyNet connect synergistic gene pairs. We followed a cross-validation procedure in which the inference of SyNet and validation of its utility in a NOP is strictly separated. We find that SyNet-based NOPs yields superior performance with higher stability across the folds compared to both the baseline model trained on all genes as well as models that use other existing gene networks. We therefore conclude that networks can improve outcome prediction, but only if this network is dataset specific and inferred from labeled training data. The Corr network, which is also dataset specific but not inferred from labeled data, also improved performance but much less than using SyNet.

A major benefit of SyNet over manually curated gene networks is that its inference is purely data driven, and therefore not biased to well-studied genes. Post-hoc interpretation of the genes selected by a NOP that utilized SyNet is therefore expected to provide a more unbiased interpretation of the important molecular players underlying breast cancer and patient survival. Analysis of the genes contained in SyNet shows strong enrichment for genes with known relevance to breast cancer. More importantly, the largest subcomponent of SyNet is strongly linked to patient prognosis because it includes many genes with a known relation to the histological grade of the tumor.

To investigate if SyNet captures known biological gene interactions, we extensively compared SyNet with existing networks. We find highly significant overlaps between links, indicating that SyNet connects genes that also have a known biological interaction. Despite this significant overlap, the majority of the SyNet links are not recapitulated by direct links in the existing networks. However, we find that accurate predictions of links in SyNet are possible if more complex graph topological descriptions of the indirect connections in the existing networks are used. Interestingly, accurate predictions are only obtained when using the breast specific networks. Apparently, although the information contained in SyNet is very similar to other gene interaction networks, they can only be exploited by the classifier if they are connected in a certain way. This might explain why using existing biological networks in NOPs directly is unsuccessful and why using graph topological measures has been successful in identifying relevant genes in outcome prediction [33,34,86].

## 4 Materials and Methods

### Inferring a synergistic network (SyNet)

We hypothesized that, in order to improve outcome prediction by network-based classification, interconnections in the network should correspond to gene pairs for which integration yields a performance beyond what is attainable by either of the individual genes (i.e. synergy). Accordingly, we formulated the synergy *S*_*ij*_ between gene *i* and gene *j* as

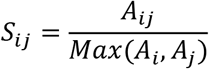

where *A*_*i*_, *A*_*j*_and *A*_*ij*_ respectively represent the Area Under Curve (AUC) of gene *i*, the AUC of gene *j* and the AUC of meta-gene *ij* formed by aggregation of gene *i* and gene *j.* Meta-gene formation is carried out by a linear regression model which demonstrated superior performance in our experiments (see **S1** for details). Cross-validation performance of the linear regression (see section on Cross validation design for details) is obtained and the median of 65 AUCs (13 folds and 5 repeats) is used as the final score *A*_*ij*_ for each pair. The AUC of the individual genes (i.e. *A_i_* and *Aj*) is obtained in a similar fashion.

Defining the synergy as a function of AUC yields a phenotype (label) specific measure which effectively ignores extraneous relationships between gene pairs that are not relevant in outcome prediction. The synergy measure *S*_*ij*_ depends on the performance of individual genes where poorly performing genes tend to achieve higher degree of synergy compared to two predictive genes (see **S2** for corresponding analysis). In order to account for this effect, the average AUC of individual genes is included as a second criterion. Furthermore, our preliminary tests confirmed previous findings [8,23,49], that integrating highly correlated genes (which reduces meta-gene noise) may improve survival prediction. For this reason, we added correlation of pairs as a third criterion. To combine these three measures, each measure is normalized and combined into an overall fitness score *F*_*ij*_ for gene pair *ij*:

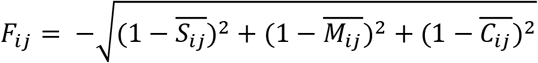

Here, *M*_*ij*_ and *C*_*ij*_, represent mean AUC and absolute spearman correlation of gene *i* and *j* respectively. Bars above letters indicate that the corresponding values are normalized to the [0, 1] interval. Employing the Dutch grid infrastructure, we quantified the fitness for all 69 million possible pairs of genes (n=11748). **Figure 1.c** visualizes the fitness of all pairs in a threedimensional space. Finally, the top 50,000 pairs with highest fitness are considered as SyNet.

### Expression Data

Accurately estimating survival risk and identifying markers relevant for progression of a complex disease such as breast cancer requires a large number of samples [11]. To this end, samples from METABRIC [50] (n=1981) are combined with 12 studies collected in ACES [21] (n=1606) as well as samples from the TCGA breast invasive carcinoma dataset [51] (n=532) (see supplementary text for details). Collectively, these datasets, spanning 14 distinct studies, form a compendium encompassing 4129 samples. To the best of our knowledge, the data combined in this paper represents the largest breast cancer gene expression compendium to date. As a result, our compendium should capture a large portion of the biological heterogeneity among breast cancer patients, as well as technical biases originating from the variability in platforms and study-specific sample preparations [52]. This variability will assist the trained models to achieve better generalization which is crucial in real world application of the final classification model [9,13,53].

To correct for technical variations that may arise during the library preparation, initially the expression data within each study is quantile normalized and then batch-effect corrected using Combat [54] where the outcome of patients was modeled as an additional covariate to maintain the variance associated with the prognostics. This procedure was shown to perform well among many batch effect removal methods [55,56]. Successful removal of batch effects were confirmed using t-SNE visualization [57] (See **S4** for details). The label for each patient corresponds to overall survival time (or recurrence free survival if available) with respect to a 5-year threshold (good vs. poor outcome).

### Regular classifiers and Network based prediction models

Ascertaining the relevance of networks in outcome prediction should be performed using a robust predictor capable of providing adequate performance in prognostic prediction. Previous assessments in this regard have been limited to only few classifiers [21,23,28,34]. To identify the optimal predictor, we have compared performance of wide range of linear and nonlinear classifiers (see **S3** for details). Supporting our previous findings [8], this evaluation demonstrates that simple linear classifiers outperform the more complex ones, with the regularized linear classifier (Lasso) reaching the highest AUC. This classifier supports both classical and well as network based prediction by its derivative called Group Lasso (GL) [58]. The GL is structurally analogous to standard Lasso with the exception of the way in which the regularization is performed; Lasso applies regularization to genes while GL enforces selection of groups of genes (See supplementary text for details). In order to incorporate network information in the GL, similar to our previous work [8], each gene in the corresponding network is considered as seed gene and together with its K neighbors the group structure provided to the GL. Priority of neighbor selection is determined by edge weights between each neighbor and corresponding seed gene. The hyperparameters for each classifier (e.g. K in the GL) are determined by means of a grid search in the inner cross validation loop (see **S5** for schematic overview).

For comparison, we include three well-known NOPs in our analysis. Park et al. utilized hierarchical clustering to group highly correlated genes [23]. Each group is summarized into a meta-gene by averaging the expression profile of the genes in that group. These meta-genes are then employed as regular features to train a Lasso classifier. The optimal cluster size for hierarchical clustering is identified by iterative application of Lasso in an inner cross-validation. Chuang et al. employs a greedy search to define subnetworks [18]. This is done by iteratively expanding a sub-network initiated from a seed gene guided by a supervised performance criterion which halts when performance no longer increases (in the training set). After groups are formed, the meta-genes are constructed by averaging expression of each gene within each group similar to Park et al. Finally, Taylor et al. focus on hubs (i.e. highly connected genes, degree>5) in a network [24]. To identify dysregulated subnetworks, the change in correlation between each hub and its direct neighbors across two classes of outcome (poor vs. good) is assessed. Meta-genes are formed from candidate subnetworks similar to the procedure employed by Park et al.

### Networks

In addition to SyNet, we considered a total of 12 other publicly available networks, including generic networks (HumanInt, BioPlex, BioGRID, IntAct and STRING), tissue specific networks [43] (brain, kidney, ovary, breast, lymph node) and a correlation network (Corr) which was previously shown to be an effective network in outcome prediction [8,23]. To the best of our knowledge, our study is the first to evaluate tissue specific networks in the context of NOPs. To maintain a reasonable network size, we utilized only the top 50,000 links (based on the link weight) in each network (similar to number of links in SyNet). For the only unweighted network, HumanInt [37], all interactions (n=~14k) were included and links were weighted according to the average degree of the two interacting genes. Moreover, a randomized version of each network is constructed by shuffling nodes in the network which destroys the biological information of the links while preserving the overall network structure (see supplementary text for full details on preparation of networks).

### Cross validation design

In order to ascertain if network information truly aids outcome prediction, the evaluation should be based on a rigorous cross-validation that closely resembles the real-world application of these models. To this end, we perform cross-study validation in order to mimic a realistic situation in which a classifier is applied to data from a different hospital than it was trained on [7]. Briefly, one study is taken out for validation of the final performance (outer loop test set). SyNet inference and NOP training are carried out on the 13 remaining studies (outer loop training set). Within each fold of the outer loop training set, again one study is left out to obtain the inner loop test set and the rest of studies for inner loop training set. The inner loop training set is sub sampled (with replacement) to 70% and regression is performed for every genes as well as gene pairs (identical set of samples are used across all genes and pairs). The AUC scores (*A*_*i*_, *A*_*j*_ and *A*_*ij*_) are calculated on the inner loop test set. This is repeated 5 times. To train a NOP for this fold, a new inner loop training set is formed by redrawing 70% of the samples from the outer loop training set. This set is also used to infer correlation network. To assess the final performance of the NOP the outer loop test set is used (see **S5** for a detailed schematic). Our initial experiments showed a large variation of performance across studies (see **S6** for details). To prevent this variation from influencing our comparisons, assignment of samples to folds in both inner and outer crossvalidation loops are kept identical across all comparisons throughout the paper. We used Area Under the ROC Curve (AUC) as the main measure of performance in this paper.

## Declarations

### Availability of data and material

The normalized and batch effect removed cohort (excluding METABRIC which requires access approval through synapse.org) can be downloaded from [87]. Furthermore, from [87], the full matrix of SyNet in binary format as well as top gene pairs (including their tri-score used to calculate their fitness) is available for download in tab-delimited format. Finally, all scripts used for preparation of data and figures in this manuscript are available for download from [88]. To ensure the full reproducibility of our results, the indices utilized for training and testing of all models (including inner and outer cross-validation) is also provided.

### Competing interests

The authors declare there are no competing interests.

### Funding

Part of computations required for this work was carried out on the Dutch national e-infrastructure (e-infra160001) with the support of the SURF Foundation.

### Authors’ contributions

AA collected, analyzed the data and prepared the manuscript. JU interpreted the results and prepared the manuscript, JdR designed the experiments, prepared and reviewed the manuscript. All authors read and approved the final manuscript.

